# Characterization of the Tau Interactome in Human Brain Reveals Isoform-Dependent Interaction with 14-3-3 Family Proteins

**DOI:** 10.1101/2022.10.17.512532

**Authors:** Ryan K. Betters, Emma Luhmann, Amy C. Gottschalk, Zhen Xu, Christopher P. Ptak, Kimberly L. Fiock, Lilliana C. Radoshevich, Marco M. Hefti

## Abstract

Tau phosphorylation and aggregation is the final common pathway for neuronal toxicity across multiple neurodegenerative diseases including Alzheimer disease, progressive supranuclear palsy, and corticobasal degeneration. We have previously shown that the fetal brain expresses high levels of phosphorylated tau, and even tau aggregates, without apparent toxic effects. The mechanisms for this remarkable resilience, however, remain unclear. In order to identify potential mediators of this resilience, we used bead-linked total tau immunoprecipitation in human fetal, adult, and Alzheimer disease brains. We then used heterologous transfection in HEK 293T cells followed by coimmunoprecipitation, mass photometry, and nuclear magnetic resonance (NMR) to further characterize the interaction of tau with one of our top hits, 14-3-3-β. We found significant differences between the tau interactome in fetal and AD brain, with little difference between adult and AD. There were significant differences in tau interaction with 14-3-3 family proteins between fetal and AD brain. We then determined that the 14-3-3 isoform with the highest difference, 14-3-3-β, preferentially interacts with 4R tau *in vitro*, forming a complex consisting of two 14-3-3-β, and one tau molecule. NMR studies using ^15^N-labeled phosphorylated tau showed that the binding site for 14-3-3 was in the microtubule binding region of tau, which is truncated in 3R tau through the exclusion of exon 10. Our findings suggest that there are marked differences between the phospho-tau interactome in fetal and Alzheimer disease brain, including differences in interaction with the critical 14-3-3 family of protein chaperones, which may explain, in part, the resilience of fetal brain to tau toxicity.

## Introduction

Aggregation of phosphorylated tau protein acts as the final common pathway for neurodegeneration across multiple neurodegenerative diseases including frontotemporal lobar degeneration with tau (FTLD-tau), chronic traumatic encephalopathy (CTE), and Alzheimer disease (AD). Although tau phosphorylation is apparently necessary, we and others have shown that it is not sufficient for tau toxicity (1,2). Human fetal tau is extensively phosphorylated, with a pattern similar to that seen in Alzheimer disease, but without apparent adverse effects (1). One of the key differences between the fetal and adult human brain is the lack of long isoform (4R) tau in the former, and this has been proposed as a potential protective mechanism (3). The remarkable resilience of human fetal brain to tau toxicity represents an unparalleled opportunity to identify mechanisms that trigger tau toxicity in Alzheimer disease and other neurodegenerative tauopathies such as corticobasal degeneration and progressive supranuclear palsy. In the current report, we present a unique data set that characterizes the tau protein interactome in fetal, adult, and Alzheimer disease brain, and identify tau splicing-dependent changes in 14-3-3 protein interaction as a potential protective mechanism in the fetal brain.

## Materials and Methods

### Human brain tissue procurement

Frozen cortical adult and fetal human brain tissue was obtained from the University of Iowa NeuroBank or the NIH NeuroBioBank. Inclusion criteria were post-mortem interval less than 24 hours and an appropriate neuropathological diagnosis (Alzheimer disease, normal control) according to published criteria (4). Exclusion criteria were gestational age less than 18 or greater than 22 post-conceptional weeks or any elective termination as defined by Iowa law. Cases with pathological evidence of global hypoxic ischemic injury were excluded. This research was reviewed by the University of Iowa’s HawkIRB and determined not constitute human subjects research under the NIH Revised Common Rule.

### Homogenization and preparation of human brain samples

Frozen brain tissue stored at −80C was mechanically pulverized using a stainless-steel mortar and pestle on dry ice. We then added 600 uL of ice-cold lysis buffer (20mM Tris, 1mM EDTA, 1mM EGTA, 240mM sucrose, 1x Halt Protease & Phosphatase Inhibitor [ThermoFisher]) to 1.5 mL tubes containing ceramic beads (#15-340-153, Fisherbrand), followed by 100 mg of pulverized frozen brain. Samples were homogenized using a Bead Mill 4 homogenizer (Fisherbrand) with velocity 5 m/s for a total of 20 seconds. The supernatant was then removed and transferred into a sterile low-protein binding microcentrifuge tube, spun at 15,000g for 30 minutes at 4 C, and the supernatant then removed. We measured final protein concentrations using a BCA (bicinchoninic acid) assay before proceeding directly to immunoprecipitation or storage at −80° C. Samples were prepared for co-immunoprecipitation through treatment with Benzonase endonuclease (#9025-65-4, Millipore) for 1ug per reaction and 15 min room temp immediately prior to co-IP.

### Brain Tissue Co-immunoprecipitation

Manual immunoprecipitation (IP) was done using mass spectrometry-compatible magnetic IP kit (#90409, ThermoFisher) with beads conjugated to an HT7 total tau antibody (#MN1000, ThermoFisher) according to the manufacturer’s directions; 500 μg of protein from each sample was combined with 5 μg of antibody and diluted to 500 μl with IP-MS Cell Lysis Buffer (#90409) before incubating overnight at 4°C with rotary mixing. Protein A/G magnetic beads were washed with IP-MS Cell Lysis Buffer in low protein binding microcentrifuge tubes and the immune-complex containing samples were then added and incubated at room temperature for one hour with constant agitation. The beads were collected using a magnetic stand and the flow-through saved for downstream analysis. Beads were then washed a total of six times for 15 minutes each with MS-grade water.

### On-bead trypsin digestion and peptide purification

Lyophilized trypsin (#V5111, Promega) was resuspended in Trypsin Resuspension Buffer (#V5111, Promega) to a concentration of 0.2 μg/μl. We then added 100μl of Trypsin Digestion Buffer (20mM Tris HCL pH 8.0, 2mM CaCl2) and 5 μl of trypsin solution to each tube. Bead-containing samples were incubated in a Thermoshaker (Eppendorf) for four hours at 37 C, shaking at 1200 rpm. The supernatant was then removed and the beads discarded. An additional 1 ug of trypsin was added to each sample followed by an additional digestion overnight for 37°C at 750 rpm on the Thermoshaker. Samples were then acidified to 1 % trifluoroacetic acid before purification with OMIX C18 pipette tips (ThermoFisher). OMIX C18 peptide purification was conducted with x5 pre-wash buffer (80% acetonitrile, 0.1% trifluoroacetic acid, Milli-Q H2O to final volume) treatments, x5 wash buffer (0.1% trifluoroacetic acid, Milli-Q H2O to final volume) treatments, and x2 elution buffer (60% acetonitrile, 0.1% trifluoroacetic acid, Milli-Q H2O to final volume) treatments. Samples were then lyophilized and stored at −80°C until needed.

### Mass spectrometry and data analysis

Purified lyophilized peptides were dissolved in 15ul loading solvent (0.1% TFA in water/CAN (98:2, v/v)) and 4uL was injected for LC-MS/MS analysis on an RSLCnano system connected to a Q Exactive HF mass spectrometer (Thermo). Trapping was done at 10uL/min for 4 min in loading solvent on a 25 mm C18 trapping column (New Objective). Peptides were then eluted using a nonlinear increase from 2 to 56% solvent B (0.1% Formic Acid in water/acetonitrile (2:8, v/v)) over 160 min at a constant flow rate of 500nL/min on a 200 cm column at 37°C. [needs more data on QExactive settings]. The resulting data were analyzed using the Andromeda search engine in MaxQuant (version 1.6.43.10) with default search settings including FDR=1% at both peptide and protein levels, mass tolerance for precursor ions at 4.5ppm, and a mass tolerance for fragment ions at 20ppm and 0.5Da. Two missed cleavages were allowed. Carbamidomethylation of cysteine residues was set as a fixed modification. Oxidation of methionine and N-terminal acetylation were set as variable modifications. Only proteins with at least one unique or razor peptide were retained in both shotgun searches. We used Perseus (version 1.6.14.0) for downstream analysis including removal of reverse database hits, potential contaminants and IDs identified only by sites. Intensities were log2 transformed and normalized for each sample by subtracting the median LFQ intensity. Replicate samples were grouped, and sites with less than three valid values in at least one group removed. Missing values were imputed from a normal distribution around the detection limit. Differential analysis was done using the Perseus package as previously described with FDR = 0.05 and S_0_=1 (5).

### HEK 293T Transfection plasmids

Plasmids for 0N3R tau, (wildtype, Ser214Ala or Ser214Asp) and 0N4R tau were the kind gift of Dr. Gloria Lee. A pcDNA3 flag HA 14-3-3 β plasmid was a gift from William Sellers (Addgene plasmid # 8999; http://n2t.net/addgene:8999; RRID:Addgene_8999). All plasmids were verified by Sanger sequencing and/or long-read sequencing (Iowa IIHG Genomics core or Plasmidsaurus, respectively). Plasmids were grown using ByLss cell transformation procedures and extracted using the EndoFree Plasmid Maxi Kit (Qiagen, 12362). Final nucleic acid concentrations were determined using a NanoDrop (Thermo, ND-2000) after washing using Qiagen tips, air drying overnight, and resuspending in endotoxin-free buffer before storing at −20°C.

### Cell culture, transfection, and lysis

HEK 293T (CRL-1573) cells were obtained from ATCC (Manassas, VA). Cells were cultured in Eagle’s Minimum Essential Medium (EMEM [ATCC, 30-2003]) with 10% heat-inactivated fetal bovine serum (Gibco, F4135). Cells were passaged every 3-5 days using Trypsin-EDTA (Gibco, 25200056). All experiments were done with cells at 10 passages or earlier with regular testing for mycoplasma. Cell transfection was done as described in (6) using a CaCl2/HeBS buffer with 5 ug total expression plasmid per 100mm dish. At the completion of the transfection protocol, cells were harvested and flash frozen as described in (7) to preserve post-translational modifications and protein-protein interactions before storing at −80°C. Cell lysates were prepared as described in (6) using modified lysis buffer (137 mM NaCl, 20 mM Tris–HCl, pH 8.0, 10% glycerol, 1 % Triton X-100) with DNase I (Millipore Sigma, DN25-10MG) and HALT (Thermo, 78446) protease and phosphatase inhibitor followed by centrifugation at 15,000 g for 30 minutes at 4°C.

### Co-immunoprecipitation and western blotting

Co-immunoprecipitation in cell lysates was performed using the same protocol as in human tissue above but using either SDS elution (2x Laemmli sample buffer (BioRab, 1610737), 5 min at 95°C) or glycine-based elution buffer (pH 2.6, 0.2M glycine•HCL, neutralized with 1-5ul of 5M NaOH prior to electrophoresis) depending on the type of downstream analysis. Gel electrophoresis and membrane transfer for Western blotting was accomplished using the Mini-PROTEAN Tetra Cell system (BioRad, 1660828EDU) with 2x Laemmli sample buffer and 2-Mercaptoethanol (BME [BioRad, 1610710XTU]). Proteins were transferred onto Immun-Blot PVDF membrane (BioRab, 1620177) and incubated overnight in blocking solution (5% powdered milk, NaF in TBST [TBS (BioRad, 1706435) 1% Tween]) or for 5 min in EveryBlot blocking buffer (Bio-Rad, #12010020) before proceeding to primary and HRP secondary antibody staining in the respective blocking buffers. TrueBlot secondary antibodies (Rockland Antibodies, 88-8887-31/88-8886-31) were used to detect the protein of interest when interference with heavy and light antibody chains or protein A/G contamination was a concern. Imaging was conducted on the BioRad ChemiDoc Imager.

### Production of ^15^N-labeled Tau441

Human Tau441 was produced in *E. coli* BL21 (DE3) cells (Novagen) using the pET29b vector (a gift from Dr. Peter Klein, Addgene plasmid # 16316) (8). Cells were grown at 37°C and 180 rpm in M9 minimal medium containing 4 g/L glucose, 1 g/L [^15^N] ammonium chloride, 1 mM MgSO_4_, 0.1 mM CaCl_2_, MEM vitamin cocktail (Sigma) and kanamycin (100 ug/mL). Non-labeled tau441 was produced using the same methods, but without ^15^N-containing growth medium. The induction was delayed until the optical density (OD_600_) reached 1.2 and initialized by addition of 0.4 mM of Isopropyl β-D-1-thiogalactopyranoside (IPTG) and continued at 37°C for 3 hours. Cells were harvested by centrifugation (5,000 g, 20 mins, 4°C) and stored at −80°C. The harvested cells were resuspended in buffer (50 mM NaPO_4_, 2 mM EDTA and 2 mM DTT, pH 6.8), and supplemented with protease inhibitor cocktail (Complete, Roche), lysozyme and DNase I. The resuspended cells were disrupted by sonication, and the cell debris was separated by centrifugation (80,000g, 45 min, 4°C). The supernatant was incubated at 75°C for 15 mins, and the soluble proteins were isolated by centrifugation (80,000g, 45 min, 4°C) and purified by cation exchange chromatography (HiTrap SP, GE Healthcare) with a gradient of 0-1M NaCl. The eluted fractions containing Tau441 were pooled and further purified by size exclusion chromatography (HiLoad 16/600 Superdex 200pg, GE Healthcare) in buffer (50 mM NaPO_4_, 2 mM EDTA and 2 mM DTT, pH 6.8). The purified protein was analyzed using SDS-PAGE and concentrated using Amicon Ultra-10K centrifugal filters.

### In vitro tau phosphorylation

We used two different *in vitro* phosphorylation techniques due to inherent differences between NMR and mass photometry and large variations in scalability of available kinase systems. For <u>large-scale</u> *in vitro* phosphorylation of ^15^N-labeled tau441 (^15^N-4R tau) bound for analysis by NMR, we utilized a two-stage process by which 3ug of activate MEK1 (MilliporeSigma, #14-429) was used to activate 50μg of ERK2 (MilliporeSigma, #14-536) prior to incubation with 4R tau. MEK1 was activated overnight at 30°C with constant agitation in phosphorylation buffer (200uL; 50mM HEPES·KOH, pH 8.0, 12.5mM MgCl2, 50mM NaCl, 1mM DTT, 1mM EGTA) with ATP (2.5mM) and HALT Protease and Phosphatase Inhibitor Cocktail (ThermoScientific, #78440) added just prior to use. We then incubated 2mg ^15^N-4R tau with 50μg activated ERK2 for 4 hours at 37°C with constant agitation (ATP was added to maintain concentration at 2.5mM despite increase in volume). The sample was then heated at 75°C for 15 minutes to inactivate the kinases prior to centrifugation at 20,000g for 15 minutes, allowing the collection of phosphorylated tau in the supernatant. The non-^15^N 4R tau used for <u>small-scale</u> chemical *in vitro* crosslinking was phosphorylated using cAMP-dependent Protein Kinase (PKA, catalytic subunit, New England Biolabs #P6000S) according to manufacturer guidelines (25uL reaction at 30°C for 2 hours with 200 μM ATP) but substituting the provided Tris-containing “NEBuffer” with an amine-free version (50mM HEPES, 10mM MgCl2, 2mM DTT, pH 7.5) to allow for unhindered downstream chemical crosslinking. PKA was inactivated by heating at 65°C for 20 minutes and cleared from the sample through centrifugation at 20,000g for 15 minutes.

### In vitro chemical crosslinking using BS^3^

We used the non-cleavable chemical crosslinker BS3 (bis(disuccinimidyl) suberate) (Thermo, #21586) according to manufacturer guidelines to perform *in vitro* amine-amine crosslinking of purified 4R tau with purified recombinant 14-3-3 β (LSBio, LS-G96740-100). Crosslinking was performed at 20-fold molar excess (5mg/uL protein concentration, 100uL reaction size) in conjugation buffer (1x PBS, pH 8.0) for 30 minutes at room temperature with gentle agitation before quenching with 1M Tris•HCL (pH 7.5) to a final concentration of 2mM with 15 minutes of incubation at room temperature. Samples that were not immediately used were stored at −80°C.

### Mass Photometry

MP experiments were performed on a Refeyn TwoMP mass photometer (Refeyn Ltd, Oxford, UK). Microscope coverslips (24 mm x 50 mm, Thorlabs Inc.) were cleaned by serial rinsing with Milli-Q water and HPLC-grade isopropanol (Sigma Aldrich) followed by drying with a filtered air stream. Silicon gaskets (Grace Bio-Labs) to hold the sample drops were cleaned in the same procedure immediately prior to measurement. All MP measurements were performed at room temperature using Dulbecco’s phosphate-buffered saline (DPBS) without calcium and magnesium (Thermo Fisher). The instrument was calibrated using a protein standard mixture: β-amylase (Sigma-Aldrich, 56, 112 and 224 kDa), and thyroglobulin (Sigma-Aldrich, 670 kDa).

Before each measurement, 15 μL of DPBS buffer was placed in the well to find focus. The focus position was searched and locked using the default droplet-dilution autofocus function after which 5 μL of protein was added and pipetted up and down to briefly mix before movie acquisition was promptly started. Movies were acquired for 60 s (6000 frames) using AcquireMP (version 2.3.0; Refeyn Ltd) using standard settings. All movies were processed, analyzed using DiscoverMP (version 2.3.0; Refeyn Ltd).

### Nuclear magnetic resonance

^15^N-labeled Tau-441 proteins were exchanged into NMR buffer (2 mM DTT, 2 mM EDTA, 50 mM NaPO_4_, pH 6.8, 10% D_2_O) and concentrated to 100 μM. ^15^N/^1^H HSQC spectra were acquired at 293 K on a Bruker AVANCE NEO 600 MHz NMR spectrometer with a gradient cryoprobe, processed using NMRPipe (9), and analyzed using POKY (10). Previously reported chemical shift assignments for ^15^N-Tau-441 (BMRB ID: 50701) (11, 12) were used to assign isolated correlation peaks. Spectra were collected for Erk2-phosphorylated ^15^N-Tau-441 alone and after addition of 14-3-3β In NMR buffer at a Tau:14-3-3β molar ratio of 1:2. Peak intensities were scaled for determination of intensity ratios.

## Results

### Co-immunoprecipitation-mass spectrometry

We first used co-immunoprecipitation mass spectrometry in post-mortem human fetal, adult and Alzheimer disease brain to identify differences in the tau interactomes between these conditions. We used bead-linked anti-total tau antibodies to immunoprecipitate tau from nine frozen samples of human frontoparietal cortex (4 fetal controls, 3 adult controls, 3 Alzheimer disease). The purpose of this study was to directly compare phosphorylated tau in fetal and AD brain, so we sought to maximize the number of fetal brains to increase the power of the comparison. Since the main purpose of this study was to compare fetal to Alzheimer disease cases, this study is deliberately underpowered for a direct comparison between adult control and Alzheimer disease brain. A total of 1093 proteins were detected, of which 234 were quantifiable. Of these, 96 differed between fetal and Alzheimer disease (**Fig 1A**), and 100 between fetal and adult control (**Fig 1B**). Interestingly, there were no significant differences in tau interactome between adult and AD brain (**Fig. 1C**). There were 75 proteins shared between the fetal-AD and fetal-adult comparisons, and 46 unique to one or the other comparison (**Fig. 1D**). A complete list of all proteins is shown in Supplemental Table 1.

**Figure 1.**
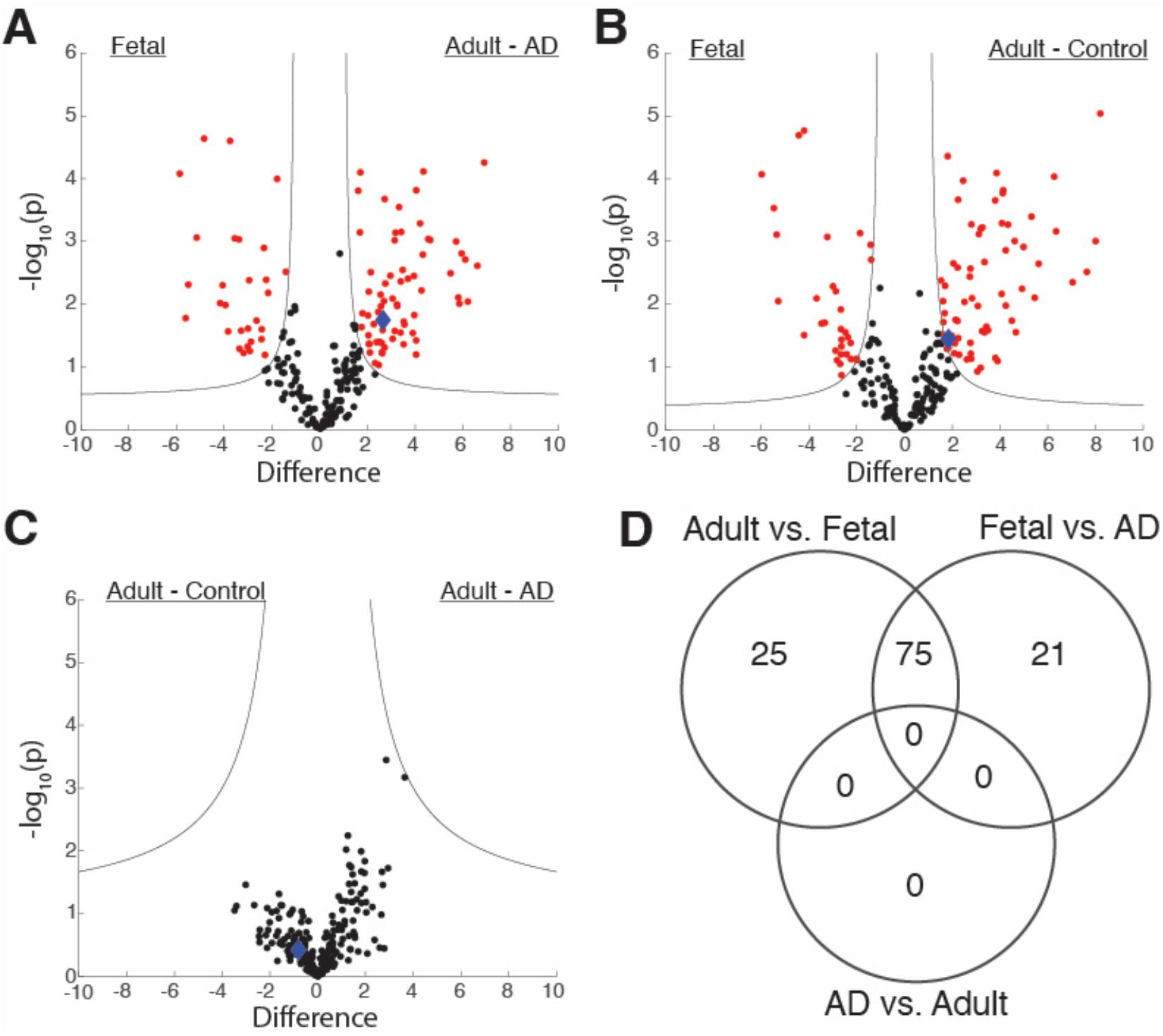
Tau interactome in human fetal, adult and Alzheimer disease brain. Volcano plot showing differentially expressed genes (red) between Fetal and AD (A), Fetal and Adult Control (B), Control and AD (C), and number of shared genes between comparisons (D).) Significant interactors are determined using a permutation-based FDR (0.05), 14-3-3-β is indicated as a blue diamond.

### Gene ontology enrichment analysis

Since both fetal and AD brain have similar levels of tau phosphorylation (1), we focused on differences between fetal and Alzheimer disease brain. We first use PantherDB to identify enriched gene ontology terms within our differentially interacting protein sets. Proteins increased in the fetal coimmunoprecipitation experiments were enriched for axon extension (GO:45773), consistent with the known role of tau as a microtubule binding protein. Interestingly, we also found enrichment for the cellular component terms paraspeckles (GO:42832), nuclear matrix (GO:16363), and nuclear speck (GO:16607). In the AD brain, the top enriched terms were neurofilament bundle assembly (GO:0033693), postsynaptic intermediate filament cytoskeleton (GO:0099160), and fructose-bisphosphate aldolase activity (GO:0004332), in the biological process, cellular component and molecular function ontologies, respectively (**Table 1**).

### Protein domain analysis

We used STRING-db to calculate enrichment of specific InterPro or PFAM protein domains in all tau interactors that could be quantified in our experiment. The top enriched InterPro domains were NOPS, fibrinogen alpha/beta/gamma chain coiled coil, microtubule associated binding protein tubulin binding, and 14-3-3 domains. Repeating the analysis using PFAM domains produced an identical result. We chose to focus on 14-3-3 family proteins for further analysis given their high level of expression in brain and known role in protein homeostasis (13).

### In vitro co-immunoprecipitation and western blotting

We then sought to validate the interaction between tau and 14-3-3 proteins, focusing on the protein with the greatest difference between fetal and AD brain (14-3-3-β). We first transfected HEK 293T cells with either 3R or 4R tau plasmids and co-transfected with FLAG-14-3-3 β. These specific isoforms were selected because, based on existing data with other 14-3-3 isoforms, the tau binding regions are thought to flank the variably spliced second microtubule binding repeat(14). We were able to co-immunoprecipitate 4R but not 3R tau with 14-3-3 β (**Fig. 2A**).

**Figure 2.**
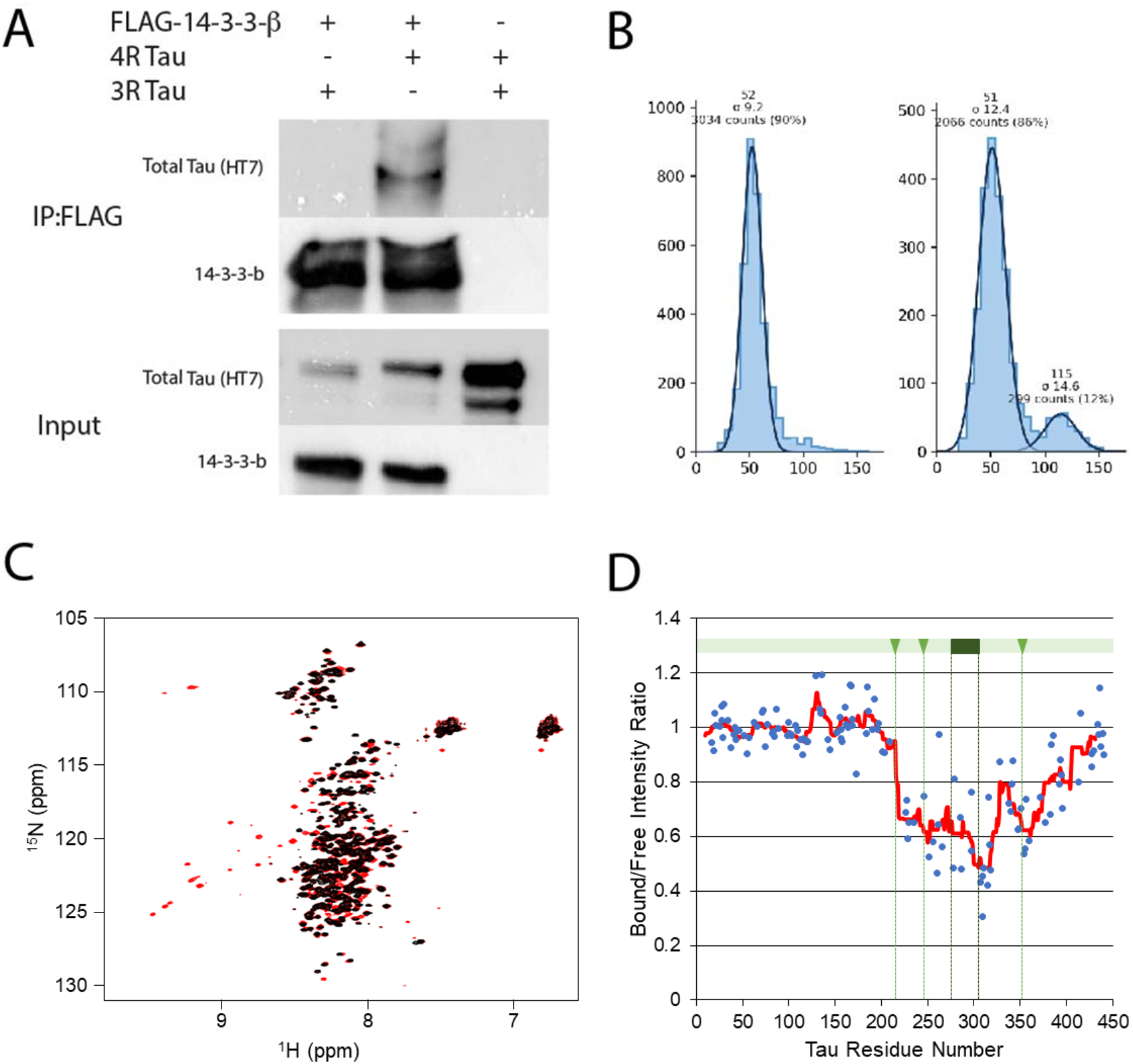
Tau −14-3-3-beta interaction depends on tau splicing and phosphorylation. (a) HEK 293T cells were transfected with the indicated constructs, immunoprecipitated with anti-FLAG antibodies, and thenb lotted with the indicated antibodies. (b) 14-3-3-β and phosphorylated (right) and unphosphorylated (left) tau were subject to mass photometry to measure individual molecule molecular weights. (c) Recombinant 15N labeled tau441 NMR with and without ERK1 phosphorylation. (d) Phosphorylated 15N-labeled tau 441 showing residue shifts after incubation with 2:1 ratio of 14-3-3-β to tau.

### Mass photometry

To further characterize this interaction, we performed *in vitro* chemical crosslinking on purified 4R (tau441) tau with and without PKA phosphorylation and 14-3-3 β and used mass photometry to measure the distribution of molecular weights in the resulting mixture. A protein complex with an approximate mass of 115 kDa was observed with phosphorylated tau, representing 12% of the total protein sample. This complex was not observed with unphosphorylated tau, which exhibited a single, 52 kDa peak consisting of both 4R tau monomers and 14-3-3 β dimers **(Fig. 2B)**. The 115 kDa species most closely aligns with a 1:2, tau:14-3-3 β complex. These data suggest a necessary role of tau phosphorylation for its interaction with 14-3-3 β and 14-3-3 proteins more generally as well as confirm 14-3-3 β’s preference for its dimerized form.

### Nuclear magnetic resonance

We then used nuclear magnetic resonance (NMR) to map 14-3-3-β interaction sites on the tau protein. Phosphorylation using ERK2 produced the expected peak shift pattern for phosphorylated amino acid residues (**Fig. 2C**). When we co-incubated phosphorylated tau with 14-3-3-β, we found a decrease in intensity for peaks corresponding to the microtubule binding region of tau (**Fig. 2D)**, overlapping with two of the three high-confidence binding sites predicted using the publicly available 14-3-3-pred algorithm. (**Fig. 2D, arrows)** (15, 16).

## Discussion

Our data shows that the greatest differences in the tau interactome are between fetal and adult brain, either control or Alzheimer disease, rather than between Alzheimer disease and adult control brain. In particular, we found that interaction between phosphorylated tau and 14-3-3 family proteins differs markedly between fetal and adult brain, and that this most likely depends on the splicing of the tau protein.

Previous studies of the tau interactome have used genetically modified tagged tau and/or animal models (17, 18). Comparing our list of quantified proteins across all age groups to that reported by Tracy et al who used APEX labeled tau in human iPSC-derived neurons, we find that 120 proteins are unique to our study, 57 are shared, and 210 are unique to Tracy et al (17). This both supports the validity of our data and underlines the importance of combining human tissue-derived data with *in vitro* studies using genetically engineered tau constructs. The limited data available in humans focuses on adult brains, either control or with neurodegenerative tauopathies and has a limited number of cases (19). None of these studies consider developmental changes or take advantage of the unique resilience of the developing brain to tau toxicity. In addition, modifying tau with either tags or with biotinylation enzymes has the potential to disrupt or change the tau interactome due to steric effects and/or the effect of tau protein overexpression. Our findings are consistent with previous data suggesting that tau interacts with 14-3-3 family proteins in both human and animal brain (20–28). Ours is however unique in that it is the first to demonstrate developmental differences in tau interaction with 14-3-3 family proteins, and describes interactions between tau and 14-3-3-β, γ, and η. 14-3-3 β has been described as a component of human neurofibrillary tangles, but its interaction with tau has not been otherwise studied (27).

Our study has several limitations. First, although we required all our cases to have short post-mortem intervals (<24 hours), we could not, due to the smaller number of specimens, include this is a covariate in our differential expression analysis for the proteomics studies. It is therefore possible, albeit unlikely, that post-mortem interval played a role in some of the observed interactions noted above. Our coIP-MS protocols are optimized to detect strong and persistent interactions – it should be noted that there are few membrane proteins in our tau interactome, suggesting that this method does not capture more transient interactions with membrane proteins. Although highly tractable and extensively used for biochemical studies, HEK 293T cells do not fully replicate the cellular environments of human neurons, and overexpressing proteins can lead to spurious interactions. The fact that we saw the same interaction, and with a consistent effect of splicing, in both our coIP-MS and transfection experiments strongly supports the validity of our data.

## Conclusions

Our data represents the first systematic characterization of the tau interactome in human fetal, adult, and Alzheimer disease brain. We report a unique data set of the human tau interactome in fetal, adult, and Alzheimer disease brain, and provide the first systematic description of tau-14-3-3 interaction in the human brain. We also present the first description of an isoform dependent tau-14-3-3-β protein interaction.

## Acknowledgements

We acknowledge the University of Iowa personnel and instrumentation in the IIHG Genomic Sequencing, the CCOM NMR, and Protein & Crystallography core facilities, supported by the Roy J. and Lucille A. Carver College of Medicine. We thank Drs. Nicholas Schnicker, Michael Dailey, and Liping Yu for helpful discussions, and wish to particularly thank Dr. Michael Dailey for his co-mentorship of the first author.

## Data Availability

The datasets supporting the conclusions of this article are included within the article and its additional file(s).

## Author Contributions

RB – Investigation, writing – initial; EL – Investigation, methodology, ACG – Formal analysis; KLF – methodology; ZX – Investigation, methodology; CP – investigation, methodology, formal analysis; LCR – methodology, supervision, formal analysis, MMH – conceptualization, supervision, writing – review and editing;.

## Classification

Biological Sciences, Neuroscience

## Ethics approval and consent to participate

Human tissue studies were reviewed by the University of Iowa’s Institutional Review Board and determined not to represent human subjects research under the NIH Revised Common Rule. All donors or their next of kin consented to the use of tissue for research in compliance with Iowa state law and all applicable U.S. federal laws and regulations. No animals were used for the reported studies.

## Consent for publication

Not applicable.

---

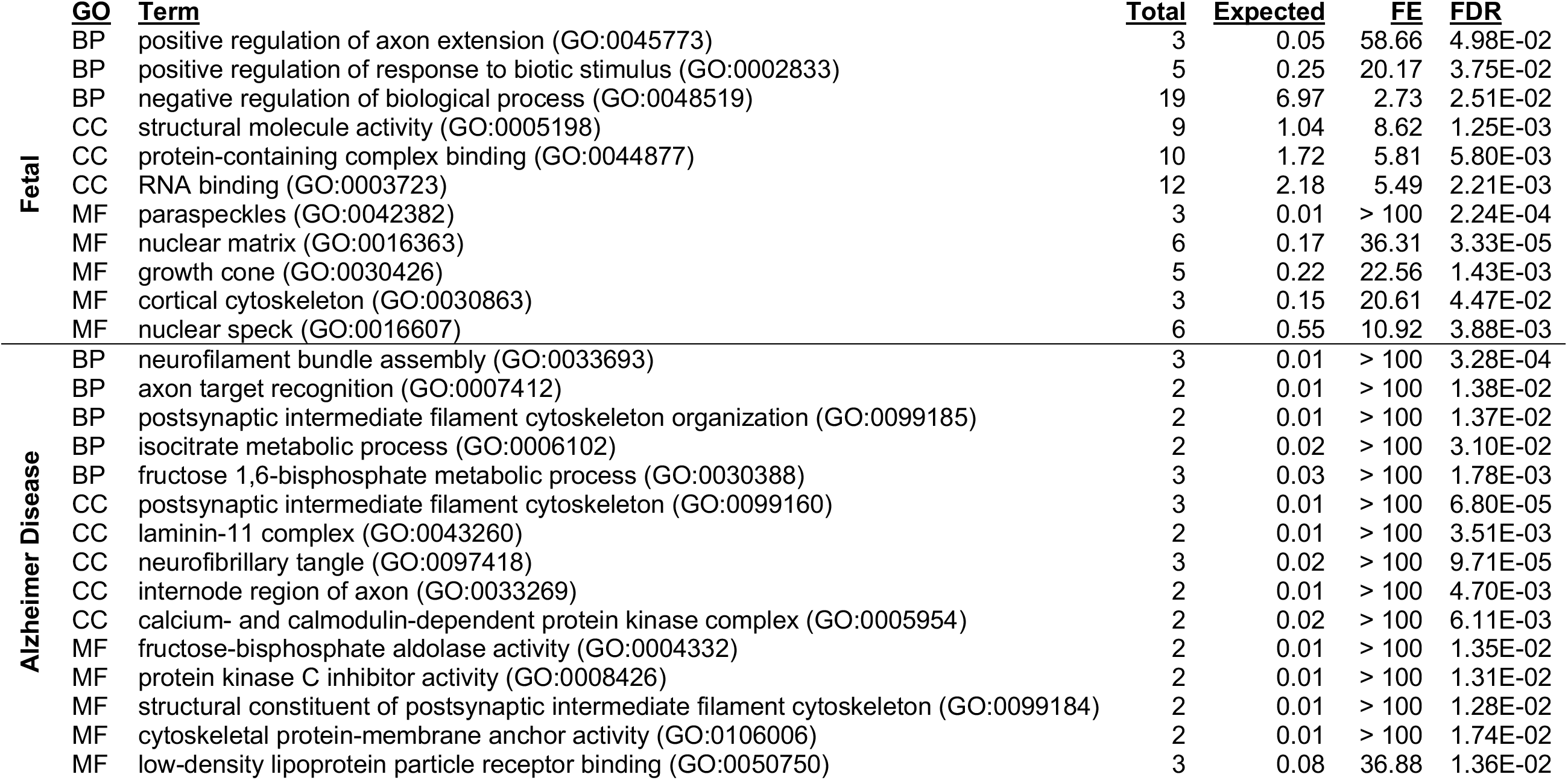

